# Distinct Cerebellar Activation Patterns Separate Novelty, Movement, and Motor Learning

**DOI:** 10.64898/2026.09.03.745918

**Authors:** Megan Sholomiski, Victor Yon, Franziska Mudlaff, Aparna Suvrathan

## Abstract

The cerebellum supports a wide range of motor and non-motor behaviors, yet the specific cerebellar regions and neuronal populations underlying this functional diversity remain poorly understood. Here, we mapped patterns of immediate early gene activation in the cerebellar cortex during the accelerating rotarod task, an established form of cerebellum-dependent motor learning. Using the immediate early gene c-Fos to identify cells active during the task, we found significantly greater activation throughout the vermis and in crus I of the hemispheres, specifically within the molecular and granule cell layers, compared to naive controls. However, a second control group, which was exposed to the rotarod apparatus but not placed on the rotating rod, revealed similar patterns of c-Fos expression, suggesting that novelty contributed substantially to the observed activation. To separate this potential novelty signal from the motor component of the task, mice were habituated to the apparatus prior to testing. This habituation resulted in an overall decrease in c-Fos expression across cerebellar lobules, with significant motor activity-related differences found in the anterior vermis. Finally, to determine how learning influences c-Fos expression patterns, mice were subjected to a multi-day motor learning paradigm in which genetic tagging of c-Fos-expressing neurons was performed on day 1, followed by c-Fos immunostaining on day 5. We found that, in mice that had learned the task, there was no significant increase in c-Fos signal above naive controls, indicating that rotarod task performance no longer recruited the same level of cerebellar cortical activity, consistent with a potentially reduced role for the cerebellar cortex once the task is well learned. Overall, these findings reveal distinct patterns of activation associated with novelty, motor behaviour, and learning, providing a framework for linking behavior to regional cerebellar function.

## 1. Introduction

The cerebellum exhibits functional heterogeneity, with studies demonstrating a diverse range of both motor and nonmotor functions (Manto et al., 2012; Strick et al., 2009). In humans, these functions have been mapped to overlapping regions across different lobules using techniques such as neuroimaging, anatomic investigations, and clinical observation of patients with stroke or lesions (King et al., 2019; Klein et al., 2016). In contrast, far less is known about the functional organization of the mouse cerebellum. Evidence from lesion, tracing, and manipulation studies suggests functional specialization, but comparison between these studies is limited by their different experimental conditions, methods, and levels of analysis. For example, in mammals, lobule IV/V of the vermis is thought to broadly support motor functions as part of the anterior network (Park et al., 2018; Sokolov et al., 2017), but evidence in mice also suggests a role in social behaviour (Chao et al., 2021) and fear (Xue et al., 2024). Conflicting evidence has been reported regarding its relationship to anxiety-like behaviors (Chao et al., 2021, 2023; Chin and Augustine, 2023).

Most of these studies use the cerebellar lobules as convenient anatomical demarcations for het-erogeneous functions, and they focus on one or a few regions of interest—a practical constraint of the techniques employed. Importantly, none provide the whole-cerebellum perspective needed to assess the relative specificity of these functions. This problem can be illustrated by comparing the functions of lobule IV/V to lobule VI−VII of the vermis. Separate mouse studies have also reported both motor (Sadakata et al., 2007) and nonmotor functions of lobule VI-VII, including cognition (Badura et al., 2018; Verpeut et al., 2023), emotion and anxiety (Ciapponi et al., 2023). However, cross-study agreement on the function of any single lobule is limited, and direct cross-lobule comparisons are rare (Chao et al., 2021, 2023). This inconsistency likely reflects identifiable methodological sources of variation: manipulation type, behavioral paradigm, and timing of perturbation can each independently determine observed outcomes (Badura et al., 2018; van der Heijden, 2024). As a result, it is not possible to determine whether the apparent functional overlap between lobules reflects genuine redundancy, shared circuitry, or an artifact of methodological heterogeneity. Yet, strong evidence for heterogeneity across different lobules/regions of the mouse cerebellum is provided by tracing studies showing heterogeneous neural connectivity (Murcia-Ramon et al., 2025; Witter and De Zeeuw, 2015b), transcriptomic studies showing molecular differences across lobules within the same cell type (Kozareva et al., 2021), and electrophysiological recordings showing differences in the baseline properties of these cells (Viet et al., 2022). These differences imply functional heterogeneity, but their relationships to behaviour-dependent activation patterns are poorly understood.

Nevertheless, mice are widely used to model human disease and disfunction, including cerebellar phenotypes of conditions such as autism (Mapelli et al., 2022), stress (Chin and Augustine, 2023), and psychiatric illness (Phillips et al., 2015; Rudolph et al., 2023). However, it remains unclear how activation patterns are distributed across the cerebellum during defined behaviors. Without a clear framework for this functional organization, it is difficult to systematically relate cerebellar manipulations in mice to specific behavioral domains or disease-relevant functions. Therefore, mapping the functional organization of the mouse cerebellum is important for the advancement of both fundamental and translational research.

Addressing this gap requires a method capable of capturing behavior-dependent activation at the level of individual lobules and cell populations across the whole cerebellum. Immediate early genes (IEGs), which are rapidly and transiently transcribed in response to cellular activation, provide a powerful tool for mapping brain activity across multiple regions. The expression of these activity markers has been studied in the cerebellum in response to various stimuli, including motor and nonmotor behaviours. However, one challenge in the comparison and interpretation of these studies is the lack of a standardized method for detecting and quantifying IEG expression in the cerebellum. Specifically, variability in how IEGs are measured, reported, and controlled for limits overall cross-study agreement. For example, due to the transient nature of IEG expression, the timing of detection is an important variable that can significantly affect experimental results (Porterfield and Mintz, 2009). Different timings have been used across published studies of induced IEG expression in the cerebellum, with no single optimized time course (Papa et al., 1993). Interpretation of these results is further complicated in some cases by a lack of detailed reporting regarding the localization of expression within particular lobules or cell populations of the cerebellum (Terao et al., 2003). Another challenge is the existence of potential confounds that are known to affect IEG expression, such as animal handling or cage changes preceding the experiment (Cho et al., 2017). These considerations, along with differences in the IEG chosen for detection (Bulthuis et al., 2025) and in the behavioural paradigm, may explain why conclusions about cerebellar IEG expression diverge even when similar motor tasks are used (Nakamura et al., 2015; Tan et al., 2024).

Therefore, careful experimentation with the appropriate controls is needed to clarify the time course, localization, and behaviour-induced activation of IEGs in the mouse cerebellum. This study aimed to address this gap using c-Fos to map patterns of activation in the mouse cerebellum following the accelerating rotarod assay, a cerebellum-dependent behaviour. The accelerating rotarod is one of the most widely used and best-validated cerebellum-dependent assays, applied across a broad range of cerebellar disease models and manipulations. We hypothesized that distinct lobules and cell populations would be activated by this task, reflecting their motor, nonmotor, and learning functions. We mapped behavior-dependent activation across all lobules of the mouse cerebellar cortex during the accelerating rotarod assay under three conditions: naive task exposure, apparatus habituation, and repeated training. Our findings reveal that the pattern of cerebellar activation is shaped by novelty and learning, providing a framework for relating cerebellar circuit manipulations to specific behavioral domains.

## 2 Results

### 2.1. The cerebellum is activated by novelty

The c-Fos fluorescence signal was analyzed separately in the granule cell layer (GCL) and molecular layer (ML) throughout the cerebellar vermis and hemispheres of wild-type mice that underwent four trials of the accelerating rotarod and compared to apparatus-only controls, which were placed at the base of the rotarod apparatus, and naive controls left undisturbed in the home cage. Mice were perfused 45 min after the final trial of the accelerating rotarod or 45 min after exposure to the apparatus based on pilot experiments demonstrating this timepoint as the approximate peak of c-Fos expression. In the hemispheres, statistically significant activation was found in the GCL (Figure 1b) and ML (Figure 1c) of Crus I in the rotarod group compared to the naive control. Interestingly, statistically significant activation was also found in the GCL and ML of apparatus-only controls compared to naive controls, and no significant separation was found between the observed signal in the rotarod and apparatus-only control groups.

In the GCL of the cerebellar vermis, statistically significant activation was found across all lobules in the rotarod group compared to naive controls, excluding the ventral sub-region of Lobule 7 (Figure 2b,d,e). This same pattern of global activation excluding the ventral sub-region of Lobule 7 was also found in the GCL of apparatus-only controls, with no statistically significant differences compared to the rotarod group. In the ML (Figure 2c,f,g), the rotarod group showed similar global patterns of activation, with a trend towards separation from the apparatus-only controls that reached statistical significance only in Lobule VI. No significant activation was found in the ML of Lobule X across any of the tested groups, likely reflecting the tonic patterns of ML activation known to support vestibular function in this lobule (Witter and De Zeeuw, 2015a). Overall, the similarity between activation patterns in the GCL and ML of the rotarod and apparatus-only control group suggests that novelty, rather than motor behaviour, may be the primary driver of this increased activation compared to naive controls.

**Figure 1.**
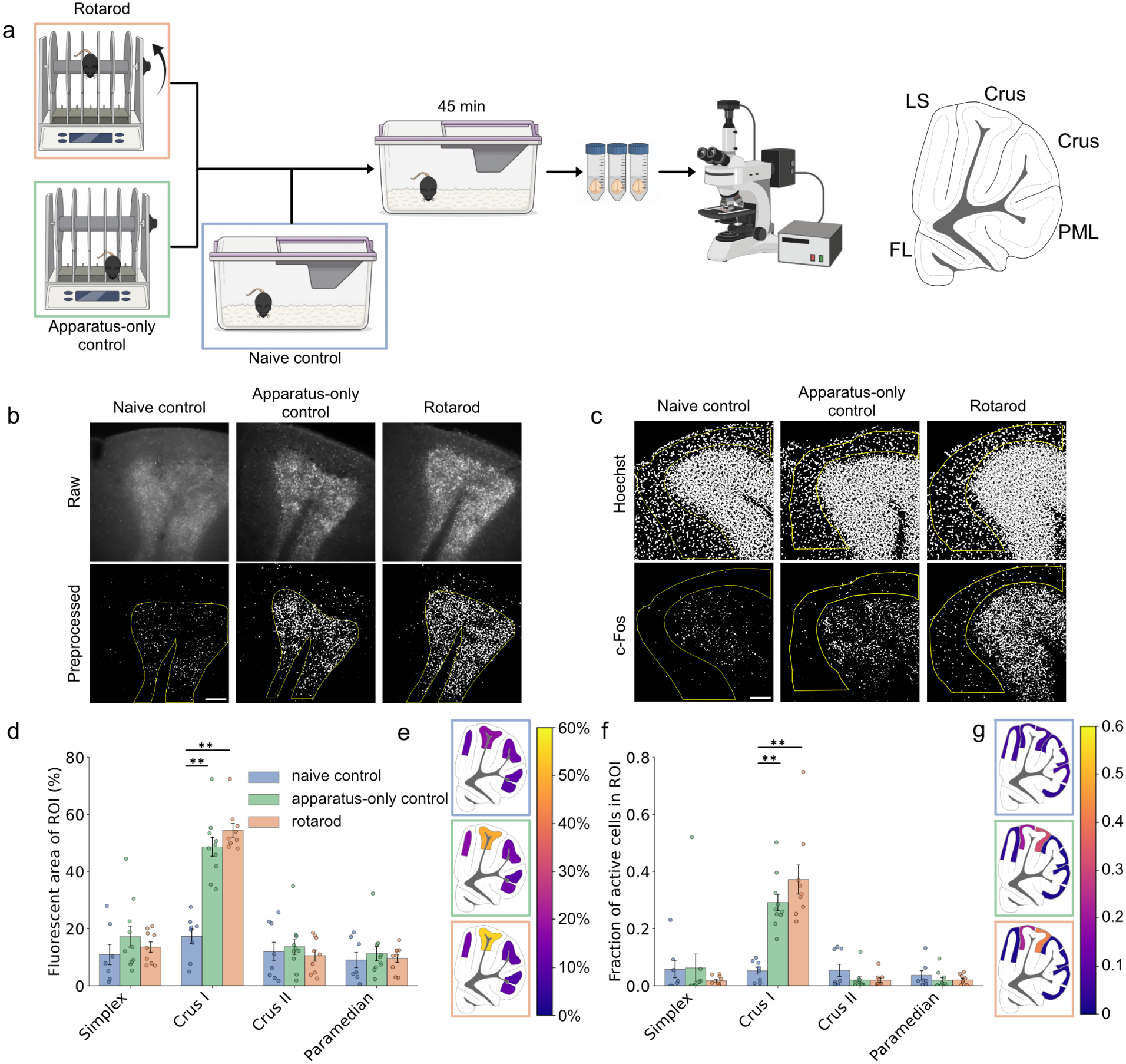
Crus I of the cerebellar hemispheres is activated by novelty. **(a)** Experimental workflow. Mice in the rotarod group (orange, *N* = 9) underwent four trials of the accelerating rotarod assay, while apparatus-only controls (green, *N* = 10) were placed on the base platform of the apparatus for an equivalent amount of time, and naive controls (blue, *N* = 8) were left undisturbed in the home cage. LS, simplex; PML, paramedian lobule. **(b)** Raw (top row) and preprocessed (bottom row) representative images of fluorescent c-Fos signal for each group in the granule cell layer of Crus I. Yellow line outlines analyzed Region of Interest (ROI). Scale bar, 100 μm. **(c)** Preprocessed representative images of fluorescent Hoechst and c-Fos signal for each group in the molecular layer of Crus I. Yellow line outlines analyzed ROI. Scale bar, 100 μm. **(d)** Bar graphs showing mean ± SEM area of c-Fos activation in the granule cell layer, normalized to the total GCL area. Each dot represents the average data of three adjacent 40 μm slices per mouse, averaged across statistically similar subregions within lobules and pooled between hemispheres (see Section 4.7). **(e)** Anatomical heatmaps corresponding to the mean data shown in (d), scaled from 0 to 60%. **(f)** Bar graphs showing mean ± SEM number of c-Fos-positive nuclei in the molecular layer of the cerebellar hemispheres, normalized to the total number of nuclei per lobule. Each dot represents the average data of three adjacent 40 μm slices per mouse, averaged across subregions within lobules that did not show statistically significant differences in activation (see Section 4.7). **(g)** Anatomical heatmaps corresponding to the mean data shown in (f), scaled as a ratio from 0 to 0.6. * *p* ≤ 0.05; ** *p* ≤ 0.01 (linear mixed model with Bonferroni correction).

**Figure 2.**
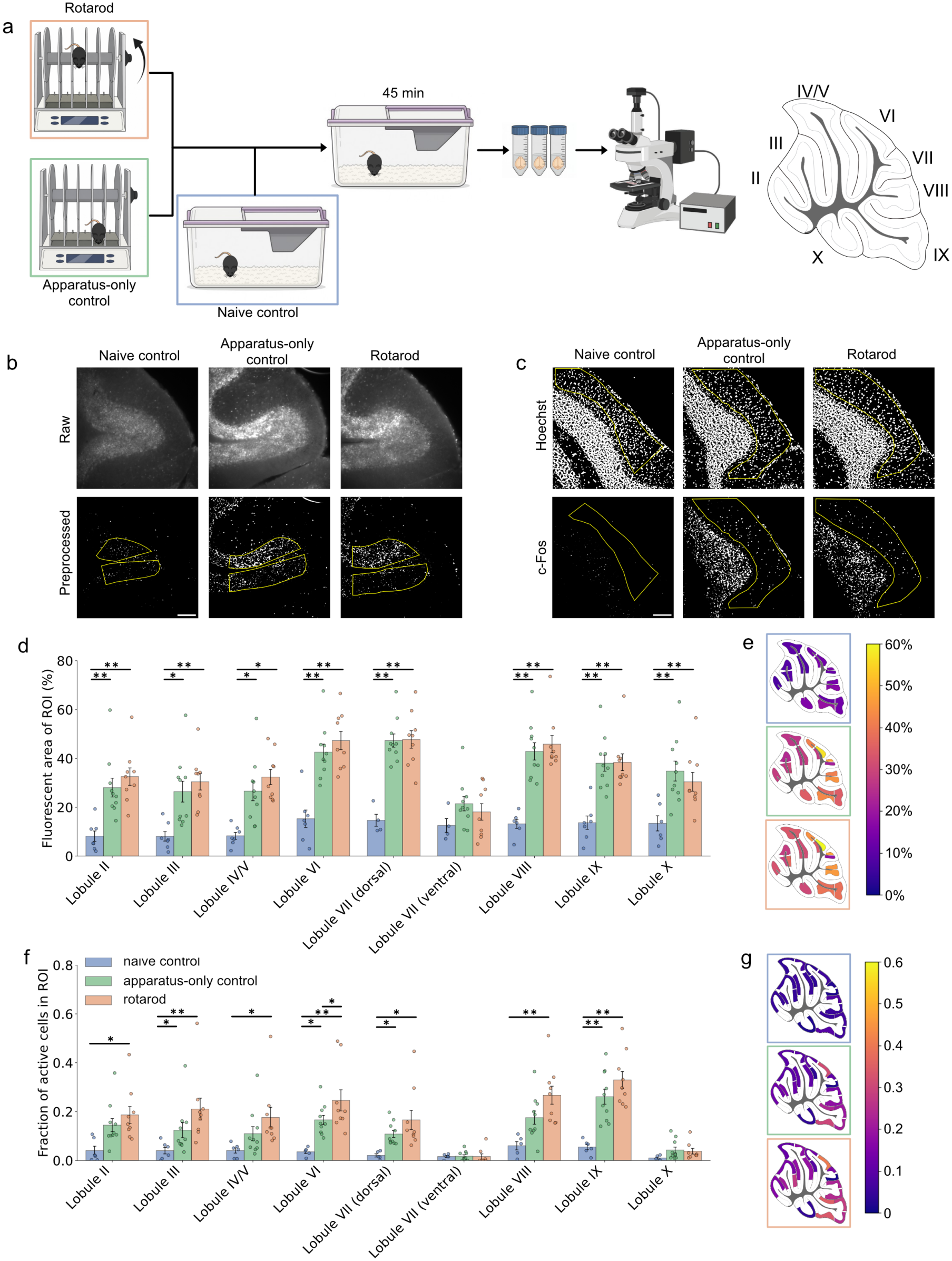
The cerebellar vermis is activated by novelty. **(a)** Experimental workflow. Mice in the rotarod group (orange, *N* = 9) underwent four trials of the accelerating rotarod assay, while apparatus-only controls (green, *N* = 10) were placed on the base platform of the apparatus for an equivalent amount of time, and naive controls (blue, *N* = 8) were left undisturbed in the home cage. **(b)** Raw (top row) and preprocessed (bottom row) representative images of fluorescent c-Fos signal for each group in the granule cell layer of Lobule VII. Scale bar, 100 μm. **(c)** Preprocessed representative images of fluorescent Hoechst and c-Fos signal for each group in the molecular layer of Lobule VI. Scale bar, 100 μm. **(d)** Bar graphs showing mean ± SEM area of c-Fos activation in the granule cell layer, normalized to the total GCL area. Each dot represents the average data of three adjacent 40 μm slices per mouse, averaged across statistically similar subregions within lobules (see Section 4.7). **(e)** Anatomical heatmaps corresponding to the mean data shown in (d), scaled from 0 to 60%. **(f)** Bar graphs showing mean ± SEM number of c-Fos-positive nuclei in the molecular layer of the cerebellar hemispheres, normalized to the total number of nuclei per lobule. Each dot represents the average data of three adjacent 40 μm slices per mouse, averaged across statistically similar subregions within lobules (see Section 4.7). **(g)** Anatomical heatmaps corresponding to the mean data shown in (f), scaled as a ratio from 0 to 60. \**p* ≤ 0.05; *p* ≤ 0.01 (linear mixed model with Bonferroni correction).

### 2.2. The anterior lobules of the vermis are activated by motor behaviour

To test whether the motor-induced activation of the accelerating rotarod assay could be separated from the putative novelty signal, the same task structure was applied to a new group of mice that were habituated to the apparatus for 3−4 days prior to the experiment (Figure 3a). Following habituation, overall c-Fos activation dropped markedly relative to the naive-exposure experiment. The apparatus-only controls no longer showed statistically significant fluorescent c-Fos signal in the molecular or granule cell layers of the hemispheres compared to the naive control group left undisturbed in the home cage (Figure 3b−g). The rotarod group showed a trend toward increased activation in Crus I, but this did not reach the level of statistical significance compared to the naive or apparatus-only controls.

**Figure 3.**
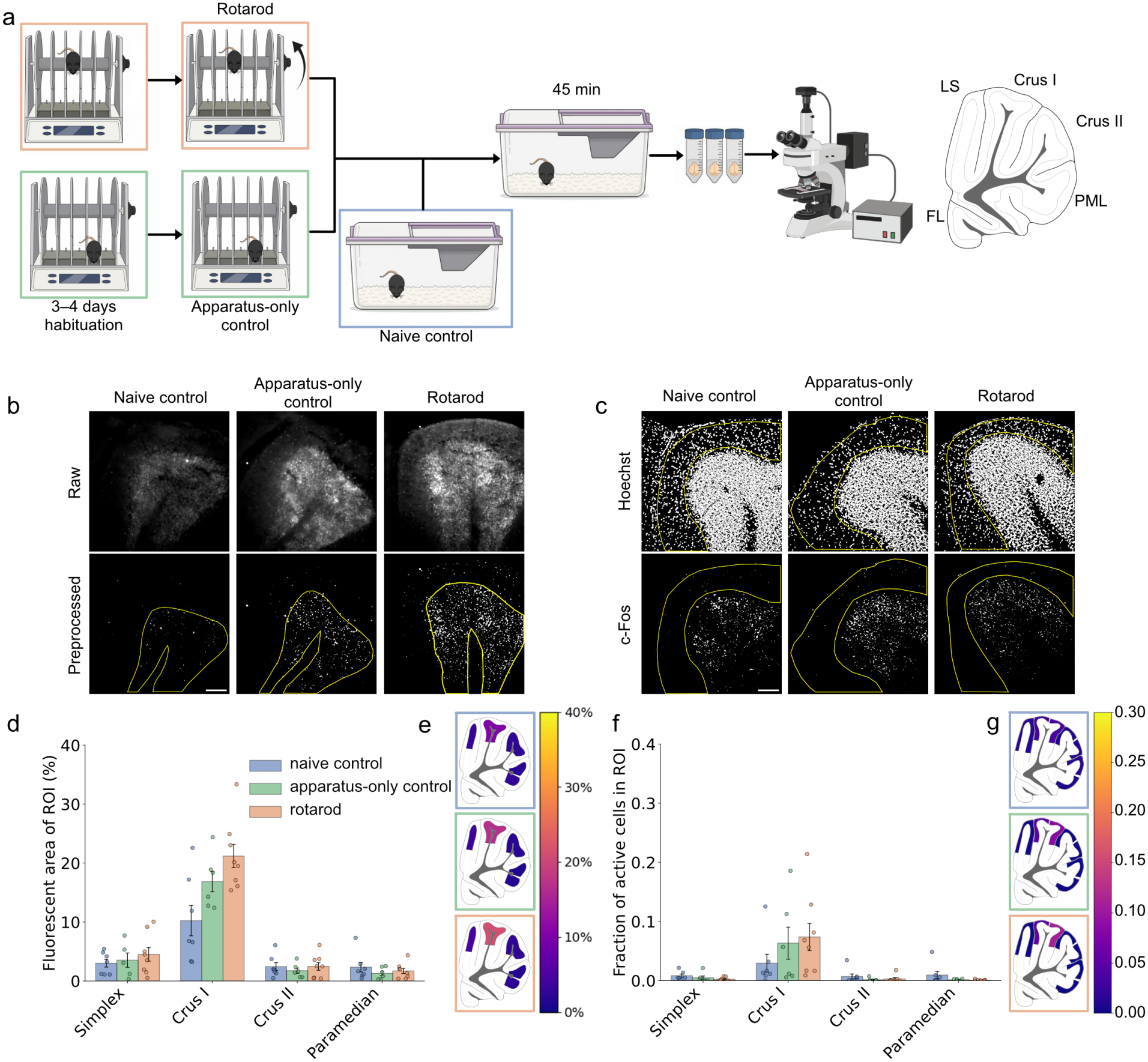
Epifluorescence imaging of c-Fos protein does not reveal significant activation of the hemispheres during the accelerating rotarod assay, **(a)** Experimental workflow. Mice in the rotarod group (orange, *N =* 8) and apparatus-only controls (green, *N* = 6) were habituated to the apparatus for 3–4 days. Then, rotarod mice underwent four trials of the accelerating rotarod assay, while apparatus-only controls were placed on the base platform of the apparatus for an equivalent amount of time, and naive controls (blue, *N = 7)* were left undisturbed in the home cage, **(b)** Raw (top row) and preprocessed (bottom row) representative images of fluorescent c-Fos signal for each group in the granule cell layer of Crus I. Scale bar, 100 pm. **(c)** Preprocessed representative images of fluorescent Hoechst and c-Fos signal for each group in the molecular-layer of Crus I. Scale bar, 100 pm. (d) Bar graphs showing mean ± SEM area of c-Fos activation in the granule cell layer, normalized to the total GCL area. Each dot represents the average data of three adjacent 40 pm slices per mouse, averaged across statistically similar subregions within lobules (see Section 4.7). **(e)** Anatomical heatmaps corresponding to the mean data shown in (d), scaled from 0 to 40%. **(f)** Bar graphs showing mean ± SEM number of c-Fos-positive nuclei in the molecular layer of the cerebellar hemispheres, normalized to the total number of nuclei per lobule. Each dot represents the average data of three adjacent 40 pm slices per mouse, averaged across statistically similar subregions within lobules (see Section 4.7). **(g)** Anatomical heatmaps corresponding to the mean data shown in (f), scaled as a ratio from 0 to 40. *\*p* ≤ 0.05; *\*\* p* ≤0.01 (linear mixed model with Bonferroni correction).

In the vermis, statistically significant activation was observed in the granule cell layer of the anterior lobules (Lobules II−IV/V) following the motor assay compared to both control groups (Figure 4d). A trend towards increased activation in the rotarod group was also observed in the GCL of Lobules VIII−IX, but this did not reach the level of statistical significance. In the molecular layer, no significant increase in c-Fos signal was observed in the rotarod or apparatus-only control groups compared to naive controls (Figure 3f). Overall, the observed separation between the rotarod and apparatus-only control groups demonstrates that habituation was effective in separating the novelty signal from motor-induced activation, reflected in the signal detected in the granule cell layer of the anterior lobules.

**Figure 4.**
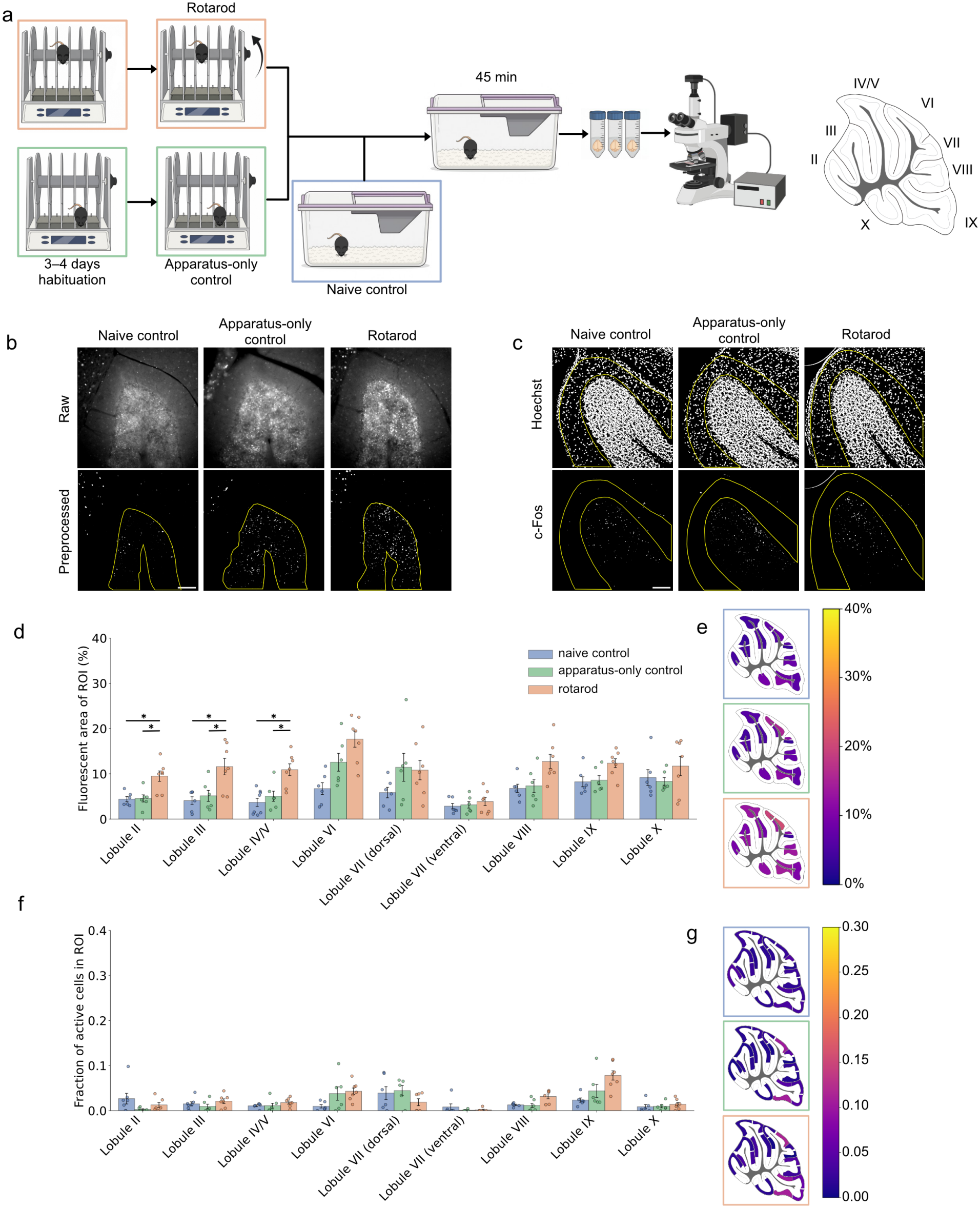
The granule cell layer of the vermis anterior lobules is activated by the accelerating rotarod assay, **(a)** Experimental workflow. Mice in the rotarod group (orange, *N = 8)* and apparatus-only controls (green, *N* = **6)** were habituated to the apparatus for 3–4 days. Then, rotarod mice underwent four trials of the accelerating rotarod assay, while apparatus-only controls were placed on the base platform of the apparatus for an equivalent amount of time, and naive controls (blue, *N = 7)* were left undisturbed in the home cage, **(b)** Raw (top row) and preprocessed (bottom row) representative images of fluorescent c-Fos signal for each group in the granule cell layer of Lobule III. Scale bar, 100 pm. (c) Preprocessed representative images of fluorescent Hoechst and c-Fos signal for each group in the molecular layer of Lobule II. Scale bar, 100 pm. **(d)** Bar graphs showing mean ± SEM area of c-Fos activation in the granule cell layer, normalized to the total GCL area. Each dot represents the average data of three adjacent 40 pm slices per mouse, averaged across statistically similar subregions within lobules (see Section 4.7). **(e)** Anatomical heatmaps corresponding to the mean data shown in (d), scaled from 0 to 40%. **(f)** Bar graphs showing mean ± SEM number of c-Fos-positive nuclei in the molecular layer of the cerebellar hemispheres, normalized to the total number of nuclei per lobule. Each dot represents the average data of three adjacent 40 pm slices per mouse, averaged across statistically similar subregions within lobules (see Section 4.7). **(g)** Anatomical heatmaps corresponding to the mean data shown in (f), scaled as a ratio from 0 to 40. *\*p* ≤ 0.05; * \**p ≤* 0.01 (linear mixed model with Bonferroni correction).

### 2.3. After mice become experts at the motor task, the cerebellar cortex is no longer activated

We next wondered whether this observed motor signal in the anterior lobules reflected a static response to balance and locomotion, or whether the signal specifically supported the early stages of motor learning, consistent with the cerebellum’s role in providing error-related feedback to motor circuits (Huvermann et al., 2025). To test whether the observed activation of the cerebellar cortex reflected motor behaviour or learning, mice were habituated to the apparatus, then re-exposed to the accelerating rotarod for five consecutive days, based on previous data from our lab showing this as the point when the learning curve plateaus. The fosTRAP2 mouse line was employed to selectively and permanently label c-Fos-positive cells within an approximately 6 h window surrounding the time of 4-OHT administration (DeNardo et al., 2019), injected immediately before the assay on Day 1 of training. This activation was subsequently compared to immunofluorescence staining of c-Fos on Day 5 to determine whether the same cell populations were re-activated after training was complete. When immunofluorescence data on Day 5 were analyzed using the same pipeline and acquisition parameters as the data presented in Sections 2.1 and 2.1, no statistically significant increase in activation was observed in the GCL or ML across any the lobules of the hemispheres (Figure 5b–e) or vermis compared to naive controls (Figure 5f–i), suggesting that when the mice had become experts at the task, the cerebellar cortex was no longer significantly recruited.

**Figure 5.**
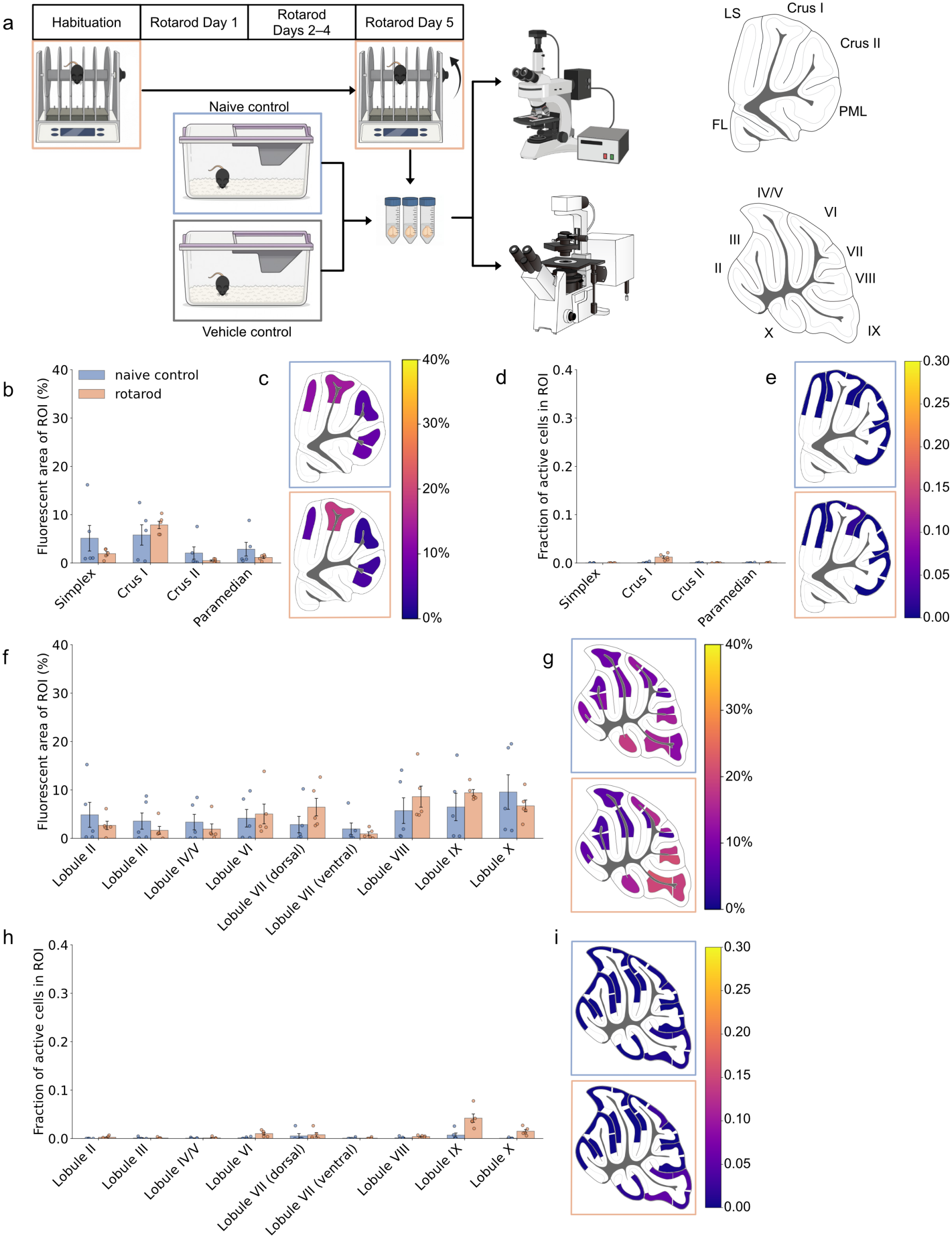
Epifluorescence imaging of c-Fos protein does not reveal significant increase in activation in the cerebellar cortex after mice have become experts at the accelerating rotarod assay, **(a)** Experimental workflow. Mice in the rotarod group (orange, *N* = 5) were habituated to the apparatus for 3–4 days, then trained on the accelerating rotarod assay for five days. 4-OHT was administered on Day 1 of training, and mice were euthanized 45 min after the final trial on Day 5 for immunofluorescence staining of c-Fos protein. Naive controls (blue, *N* = 5) received 4-OHT injections and vehicle controls (grey, *N* = 5) received vehicle injections, then both groups were sacrificed for immunofluorescence staining of c-Fos protein five days later, **(b)** Bar graphs showing mean ± SEM area of c-Fos activation in the granule cell layer of the cerebellar hemispheres, normalized to the total GCL area. Each dot represents the average data of three adjacent 40 pm slices per mouse, averaged across statistically similar subregions within lobules (see Section 4.7). **(c)** Anatomical heatmaps corresponding to the mean data shown in (b), scaled from 0 to 40%. **(d)** Bar graphs showing mean ± SEM number of of c-Fos-positive nuclei in the molecular layer of the cerebellar hemispheres, normalized to the total number of detected nuclei and scaled as a ratio from 0 to 40. Each dot represents the average data of three adjacent 40 pm slices per mouse, averaged across statistically similar subregions within lobules (see Section 4.7). **(e)** Anatomical heatmaps corresponding to the mean data shown in (d), scaled as a ratio from 0 to 40. **(f)** Bar graphs showing mean ± SEM number of c-Fos-positive nuclei in the granule cell layer of the cerebellar vermis, normalized to the total GCL area. Each dot represents the average data of three adjacent 40 pm slices per mouse, averaged across statistically similar subregions within lobules (see Section 4.7). **(g)** Anatomical heatmaps corresponding to the mean data shown in (f) scaled as a ratio from 0 to 40. (h) Bar graphs showing mean ± SEM number of of c-Fos-positive nuclei in the molecular layer of the vermis, normalized to the total number of detected nuclei and scaled as a ratio from 0 to 40. Each dot represents the average data of three adjacent 40 pm slices per mouse, averaged across statistically similar subregions within lobules (see Section 4.7). **(i)** Anatomical heatmaps corresponding to the mean data shown in (h), scaled as a ratio from 0 to 40. *\*p* ≤0.05; *\*\* p ≤*0.01 (linear mixed model with Bonferroni correction).

Confocal stacks of select lobules of the cerebellar vermis and hemispheres were acquired to compare fosTRAP labeling on Day 1 to immunofluorescence staining on Day 5 in the granule cell layer at a single-cell resolution. For fosTRAP labeling across the acquired lobules, no statistically significant differences in cell counts were found between the naive control and rotarod groups, or between the vehicle and naive control groups. There was a statistically significant separation between the rotarod and vehicle control groups in Crus I, Crus II, and Lobules II, VI, VIIv, and IX (Figure 6c). When fosTRAP (Day 1) and IF (Day 5) active cell counts were compared between the rotarod and control groups, no statistically significant differences were detected (Figure 6d), indicating that re-activation in the experimental group did not surpass chance levels (Figure 6e). These results further support that when mice become experts at the accelerating rotarod assay, previously active populations in the GCL are no longer significantly recruited.

**Figure 6.**
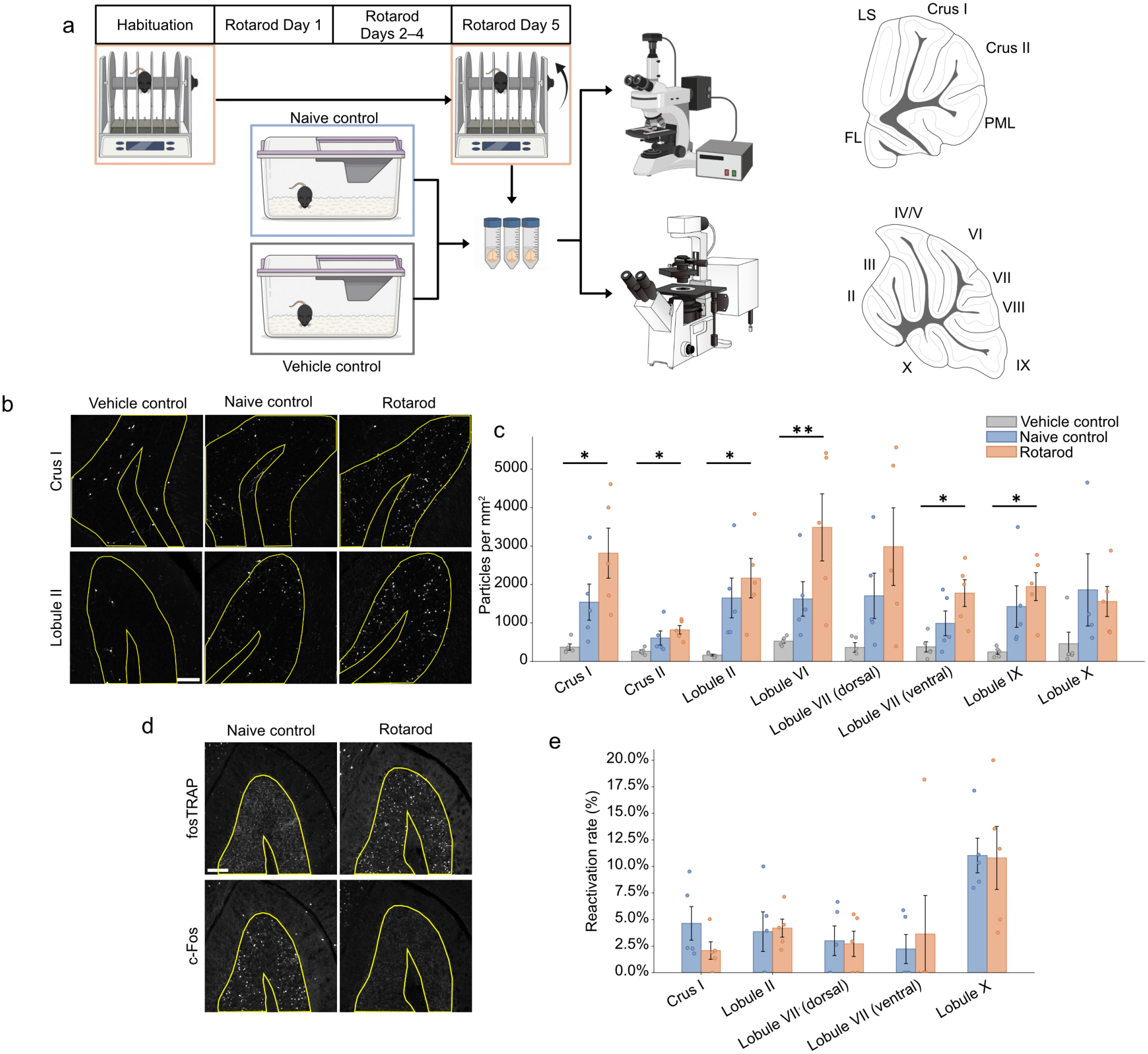
The granule cell layer is not re-activated above chance levels after mice learn the accelerating rotarod assay. **(a)** Experimental workflow. Mice in the rotarod group (orange, *N* = 5) were habituated to the apparatus for 3 –4 days, then trained on the accelerating rotarod assay for five days. 4-OHT was administered on Day 1 of training, and mice were euthanized 45 min after the final trial on Day 5 for immunofluorescence staining of c-Fos protein. Naive controls (blue, *N* = 5) received 4-OHT injections and vehicle controls (grey, *N* =5) received vehicle injections, then both groups were sacrificed for immunofluorescence staining of c-Fos protein five days later. **(b)** Representative confocal images (maximum intensity z-projections) of TdTomato reporter in fosTRAP+ cells in the vehicle control (left), naive control (center), and rotarod (right) groups. Yellow outline shows analysed area surrounding the granule cell layer of Crus I (top row) and Lobule II (bottom row). Scale bar, 100 µm. **(c)** Bar graph showing mean *±* SEM total number of fosTRAP+ cells within GCL, normalized to total ROI area (grey, vehicle control; blue, naive control; orange, rotarod). **(d)** Representative confocal images (maximum intensity z-projections) of TdTomato reporter in fosTRAP+ cells (top) and c-Fos immunofluorescence (bottom) of Lobule III in the naive control (left), and rotarod (right) groups. Scale bar, 100 µm. **(e)** Bar graph showing average fraction of fosTRAP+ cells (Day 1 activation) that are also c-Fos+ (Day 5 activation) based on immunofluorescence staining. \**p≤* 0.05; * \**p≤* 0.01 (linear mixed model with Bonferroni correction).

## 3. Discussion

Immunofluorescence staining for c-Fos protein was used in three experimental paradigms to separate the contributions of exposure to a novel environment, motor behaviour, and learning throughout the entire cerebellar cortex during the accelerating rotarod assay. First exposure to the assay produced widespread activation of the vermis and of Crus I of the hemispheres that did not differ between the rotarod and apparatus-only control groups, implicating novelty rather than motor engagement as the dominant driver of the naive-state signal. The role of novelty in activating the cerebellum was validated by the observation that habituation to the apparatus abolished this novelty signal and revealed a residual, smaller, motor-related activation in the granule cell layer of the anterior vermis. Once animals were trained to a behavioural plateau, cortical activation was no longer detectable, and neurons active on the first day of training were not reactivated above chance once the task had been learned. Together, these results provide a whole-cerebellum, layer-resolved map that dissociates novelty, motor, and learning components of a canonical cerebellum-dependent behaviour.

The similarity between the rotarod and apparatus-only groups, together with the loss of significant apparatus-only activation after habituation, indicates that the activation observed in naive animals reflects novelty rather than the motor demands of the task. A comparable pattern has been reported previously: Nakamura et al. (2015) found robust c-Fos induction throughout the vermis and in Crus I following the rotarod in wild-type mice, and they further showed that handling and exposure to a stationary apparatus were each sufficient to induce cerebellar c-Fos. Our apparatus-only control formalizes this observation and shows that, without habituation, novelty-driven activation is indistinguishable from task-driven activation across most of the cerebellar cortex. This carries a direct methodological implication: studies that map cerebellum-dependent behaviour without a novelty or apparatus control risk attributing novelty-related activation to the behaviour under study. Uncontrolled novelty of this kind is a plausible contributor to the inconsistent activation patterns reported across previous immediate early gene studies of the cerebellum (Cho et al., 2017; Porterfield and Mintz, 2009).

The localization of hemispheric activation to Crus I is consistent with the associative, rather than motor, function of this region. In humans, Crus I/II contributes to the executive-control and default-mode networks (Buckner et al., 2011; Habas et al., 2009; King et al., 2019), and in mice, manipulation of Crus I disrupts social and cognitive behaviour without impairing motor coordination (Badura et al., 2018; Stoodley et al., 2017). Activation of Crus I during first exposure to the apparatus is therefore more parsimoniously explained by the processing of a novel context than by motor engagement. Consistent with this, naive control animals that were euthanized during the active phase of the circadian cycle (when locomotor activity is highest) showed little Crus I signal, indicating that the activation seen in the experimental groups was not a general consequence of movement or arousal.

In the vermis, activation was found across the molecular and granule cell layers of all lobules, with two exceptions: the ventral part of Lobule VII, and the molecular layer of Lobule X. This observation held for both the unhabituated rotarod and apparatus-only control groups. The absence of molecular layer activation in lobule X is consistent with the tonic, vestibular-related activity of this region (Barmack, 2003; Witter and De Zeeuw, 2015a); because activity in the vestibulocerebellum is high at baseline, task-related increases may be difficult to resolve against this background. Therefore, the difference between the dorsal and ventral subregions of lobule VII is the clearest evidence in this dataset that behaviour-dependent activation is not uniform within a lobule.

Activation in the molecular layer broadly mirrored the patterns observed in the granule cell layer across lobules, although this relationship was less apparent for the residual motor-related signal revealed after habituation. This correspondence between the granule cell layer and the molecular layer is consistent with the recruitment of the well-described excitatory pathway from granule cells to molecular layer interneurons (Mittmann et al., 2005). Although we did not directly measure Purkinje cell activity, these findings raise the possibility that granule cell activation is accompanied by changes in feedforward disynaptic inhibition of Purkinje cells. Thus, while the present data do not provide a direct readout of cerebellar cortical output, they suggest that activity propagates beyond the granule cell layer to engage downstream elements of the cortical microcircuit.

One interpretation of these finding could be that the cerebellar cortex is organized into parasagittal zones and molecular compartments that cross lobule boundaries (Apps and Hawkes, 2009; Apps et al., 2018; Cerminara et al., 2015). The prototypical marker of this organization is zebrin II (aldolase C), whose reproducible banding correlates with differences in Purkinje cell firing (Xiao et al., 2014; Zhou et al., 2014) and with broader transcriptomic heterogeneity within cell types across lobules (Kozareva et al., 2021). However, parasagittal molecular zones subdivide the cortex along the mediolateral axis and are present in all lobules (Apps and Hawkes, 2009; Cerminara et al., 2015), and band-selective recruitment would be expected to pattern activation across the vermis rather than a discrete subregional difference within Lobule VII. A within-lobule anterior-posterior functional gradient is similarly insufficient, as it does not account for the comparable activation of the other posterior lobules (Stoodley and Schmahmann, 2009). Further experimentation is required to determine the functional basis of this dorsal-ventral difference in Lobule VII; however, this finding holds conceptual significance: activation that divides a single lobule indicates that the lobule is too coarse a unit for functional attribution, consistent with the view of the cerebellar cortex as an assembly of non-uniform microcircuits rather than a set of functionally homogeneous lobules (Cerminara et al., 2015).

Following habituation, which eliminated the novelty signal, significant task-related activation remained only in the granule cell layer of the anterior vermis (Lobules II–IV/V). This localization is consistent with the classical association of the anterior vermis and spinocerebellum with motor control (Manni and Petrosini, 2004; Stoodley and Schmahmann, 2009) and demonstrates that a motor-related signal can be isolated from novelty once the latter is controlled by habituation. In the hemispheres, the Crus I activation seen in naive animals fell to a non-significant trend after habituation, confirming that novelty accounted for the majority of the earlier hemispheric signal. In the posterior lobules (VIH–IX), there was a trend towards increased activation in the rotarod group compared to the apparatus-only and naive controls, but this did not reach the level of statistical significance.

When animals were trained to a behavioural plateau, no significant increase in c-Fos activation was detected in the granule or molecular layers of any lobule, in either the vermis or the hemispheres, relative to naive controls. At a single-cell resolution, neurons labeled on the first day of training by fosTRAP were not reactivated above chance in the trained state. The disappearance of the cortical signal once the task was learned is consistent with a role for the cerebellar cortex in the acquisition, rather than the expression, of the learned behaviour. During early learning, the cortex receives error-related signals that drive plasticity (Kimpo et al., 2014), whereas the consolidated behaviour may be expressed through downstream circuits, including the deep cerebellar nuclei and their brainstem and thalamo-cortical targets (Lisberger, 2021; Varani et al., 2026). Under this interpretation, the early motor signal in the anterior vermis is relevant for ongoing learning in the cerebellar circuit rather merely reflecting input signals related to locomotion and balance.

Certain limitations qualify these conclusions. First, the absence of a detectable increase in c-Fos cannot be equated with the absence of neural activation. In this context, noise in immunostaining, threshold sensitivity, and a potential epifluorescence detection floor could all contribute to the potential loss of small but biologically significant differences in c-Fos expression. However, whole-cerebellum, layer-resolved mapping requires uniform application across the entire cortex, which higher-resolution methods cannot provide at this scale. Therefore, we selected this method despite its known limitations, applying conservative preprocessing and thresholding parameters to limit false positives and interpreting results as relative rather than absolute.

IEGs are also known to report only a subset of active neurons (Kovacs, 2008), and expression differs by cell type. Purkinje cells (PCs) are tonically active and have been shown to undergo both long-term depression and potentiation in response to learning (Coesmans et al., 2004; Gao et al., 2012; Raman and Bean, 1999), confounding a straightforward readout of c-Fos as an activity marker. Our analysis therefore focused on the molecular and granule cell layers, providing a readout of activation throughout the vermis and hemispheres but omitting the cortical output layer. We do not infer Purkinje cell recruitment from granule cell layer activation: although the ascending granule cell axon provides spatially restricted input to a small number of overlying Purkinje cells, the parallel fibre system distributes granule cell output across hundreds of Purkinje cells and multiple microzones (Valera et al., 2016), and the relative functional weight of these two pathways remains unresolved. Granule cell layer activation is thus best interpreted as an index of cortical input and local processing rather than of cortical output, and direct measurement of Purkinje cell activity would be required to complete the circuit-level map.

The fosTRAP analysis carries an additional caveat: fosTRAP expression on the first day of training separated the rotarod group from vehicle controls but not from naive controls, so the baseline against which reactivation was assessed was itself weakly defined, limiting the strength of the reactivation conclusion. This result contrasts with the immunofluorescence data from the same stage of learning, in which rotarod training produced significant activation of the anterior vermis relative to both control groups. Several methodological differences could account for this mismatch. Unlike immunofluorescence, which provides a snapshot of c-Fos protein expression at a defined timepoint, fosTRAP labeling depends on CreERT2-mediated recombination and subsequent reporter expression exceeding a detection threshold. Furthermore, fosTRAP integrates activity over an approximately 6 h window following 4-OHT administration (DeNardo et al., 2019) rather than the shorter time window captured by immunofluorescence. A relatively small task-related signal may therefore be detectable using immunofluorescence yet become diluted when integrated across several hours. Consequently, the Day 1 fosTRAP signal may have lacked the sensitivity required to reproduce the group separation observed using immunofluorescence. Importantly, the lack of separation between the rotarod and naive groups in the fostTRAP experiment does not necessarily indicate the absence of learning-related activation on Day 1 but rather that fosTRAP and immunofluorescence approaches produced different estimates of that activation. The interpretation of this result as consolidation of learning-related circuits rests on an absence of detectable signal; therefore, this account remains a hypothesis that should be observable as a shift of activation from cortex to the cerebellar nuclei and/or their brainstem and thalamo-cortical targets (Lisberger, 2021; Varani et al., 2026).

Finally, our experimental design is correlational and rests on a single detection timepoint and behavioural paradigm, whose specific motor demands may not generalize to motor learning more broadly. The correlational nature of this map calls for causal interrogation: activating or silencing the anterior vermis granule cell population during early training would establish whether this activation is required for acquisition rather than merely coincident with it. And because the map rests on a single behaviour, extending it to other cerebellum-dependent paradigms would separate what is specific to the accelerating rotarod from what generalizes.

Overall, these findings provide a whole-cerebellum, layer-resolved map of behaviour-dependent activation across the lobules of the cerebellar cortex during the accelerating rotarod assay, and they dissociate the contributions of novelty, motor behaviour, and learning to that activation. The identification of novelty as a dominant driver of cerebellar IEG expression establishes habituation and apparatus controls as necessary components of cerebellar activity-mapping studies. The restriction of the motor-related signal to the anterior vermis, and its disappearance once the task is learned, motivate direct interrogation of downstream targets to test whether the learned behaviour is expressed through these circuits. More broadly, this framework provides a basis for relating cerebellar circuit manipulations in mice to defined behavioural domains, of relevance both to fundamental cerebellar physiology and to the modelling of cerebellar contributions to disease.

## 4. Methods

### 4.1. Mice

Male and female adult (3–6 months) C57BL/6J mice and Fos^tm2.1(icre/ERT2)Luo^/J (TRAP2) mice were used for immunofluorescence experiments. To evaluate learning-induced changes in c-fos expression, Fos^CreERT2^ mice were crossed with B6.Cg-Gt(ROSA)26Sor^tm14(CAG-tdTomato)Hze/J^ (Ai14) reporter mice. This cross enables 4-hydroxytamoxifen-inducible expression of the tdTomato fluorescent reporter in c-fos-positive cells. All mice were housed in standard individually ventilated cages on a 12 h reverse light/dark cycle with ad libitum access to water and standard rodent diet. All experiments were performed in accordance with the policies of the Canadian Council on Animal Care, using protocols approved by the Montreal General Hospital Facility Animal Care Committee.

### 4.2. Accelerating Rotarod Assay

A standard method for the accelerating rotarod assay was used in which mice were placed on a rotarod (Harvard Apparatus), moving at 4 RPM. After the mice were placed on the rotarod, the bar accelerated from 4 to 40 RPM over 5 min, then remained at 40 RPM for up to 5 min until the mice fell. This was repeated four times, with 10 min intervals between trials. For the rotarod control group, mice were placed on the base platform of the rotarod for an equal amount of time.

### 4.3. Tissue Preparation

Mice were perfused with 1× phosphate-buffered saline (PBS) followed by 4% paraformaldehyde (PFA) 45 min after the final trial of the accelerating rotarod assay for immunofluorescent staining of c-Fos. This timing was chosen based on pilot experiments performed to determine the peak c-Fos signal in the cerebellum following the behavioral paradigm. The brains were extracted and kept in PFA overnight at 4 °C, then transferred to 30% sucrose in 1× PBS with 0.05% sodium azide. The brains were considered equilibrated when they sank to the bottom of the tube, then the cortex and cerebellum were separated and embedded in optimal cutting temperature (OCT) compound, frozen, and stored at −80 °C. Prior to sectioning, the brains were transferred to −20 °C, then cut into 40 μm sagittal sections using a cryostat. Sections from the cerebellar hemispheres and vermis were collected and placed into 24-well plates for immunofluorescent staining.

### 4.4. Immunofluorescence Staining

Brain sections were washed three times for 10 min in PBST (0.3% Triton-X 100, Fisher Scientific, BP151-100), blocked for 1 h in 10% normal goat serum (Gibco™, 16210064) in PBST, then washed three times. Polyclonal antibodies for c-fos (Synaptic Systems, 226 008) and calbindin (Invitrogen, PIPA5143561) were diluted 1:2000 in PBST, then applied overnight at 4 °C. The following day, the sections were washed three times, then fluorescent secondary antibodies (C57/Bl6: goat anti-rabbit IgG AlexaFluor™ 568 Invitrogen, A-11011; goat anti-chicken IgY AlexaFluor™ 647, Invitrogen, A-21449. fosTRAP: goat anti-rabbit IgG AlexaFluor™ 617, Invitrogen, A-21245; goat anti-chicken IgY AlexaFluor™ 488, A-11039) were diluted 1:500 in PBST and applied for 2 h at room temperature with Hoechst 33342 (1:10,000, Invitrogen, 62249). Finally, the sections were washed three times and mounted onto polarized microscope slides (Fisherbrand, 12-550-15) with Fluorshield mounting medium (Sigma-Aldrich, F6182). The slides were imaged at 4× and 20× magnification using an Olympus BX61 epifluorescent microscope to acquire images of the entire cerebellar cortex (vermis and hemispheres) and an Olympus BX61 confocal microscope to acquire z-stacks of select lobules.

### 4.5. Tamoxifen-Inducible Reporter Activation

4-Hydroxytamoxifen (4-OHT) (Sigma-Aldrich, H6278) was dissolved in 100% ethanol at 60 °C to a concentration of 100 mg/mL, aliquoted, and stored at −20 °C. On the day of the experiment, frozen aliquots were re-dissolved at 60 °C, then diluted 1:10 in a 1:4 mixture of castor oil and sunflower seed oil to obtain a final concentration of 10 mg/mL. Immediately before the first trial of the accelerating rotarod assay (rotarod group), 50 mg/kg of the dissolved 4-OHT was injected intraperitoneally into each mouse. After the assay was complete (four trials total), the mice were returned to their home cages and left undisturbed until the following day of training. For the naive control group, mice received injections of 4-OHT, then were immediately returned to the home cage and left undisturbed for five days before being perfused for IF staining. A second control group received injections of vehicle without 4-OHT, were immediately returned to the home cage, and were sacrificed five days later. TdTom reporter expression in select cerebellar lobules was imaged using an Olympus BX61 confocal microscope.

### 4.6. Fluorescence Quantification Pipeline

Separate regions of interest (ROIs) were drawn in ImageJ (2.14.0) surrounding the complete granule cell layer (GCL) and molecular layer (ML) in each acquired image, excluding areas of damage or debris obstructing the view of the tissue. All subsequent analysis was performed in Python (3.12). A schematic and description of the pipeline is provided below (Figure 7), and the complete analysis script is available on GitHub at: https://github.com/victor-yon/fluorescent-area-analysis.

**Figure 7.**
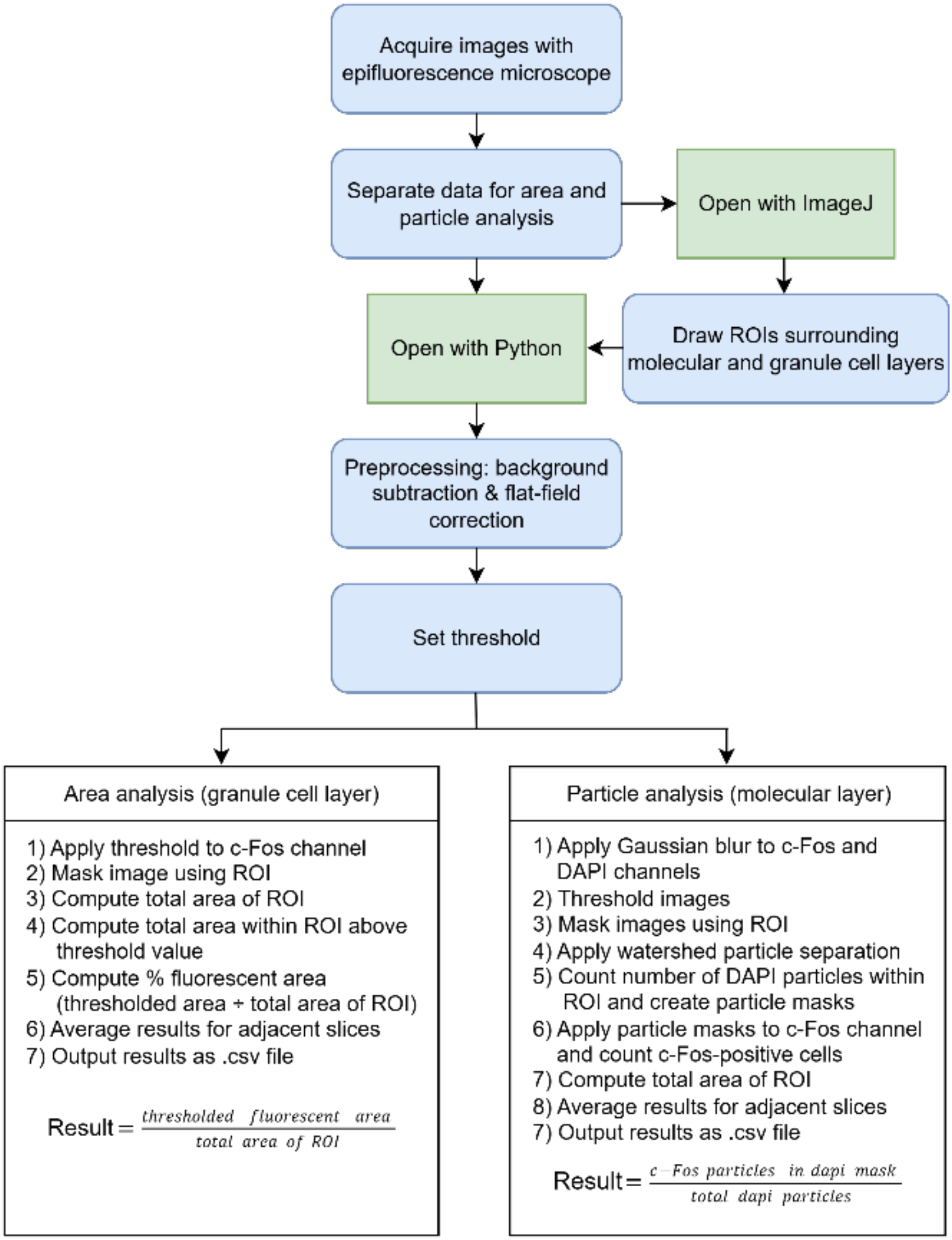
Schematic of the automated analysis pipeline used for epifluorescence image processing.

#### 4.6.1. Image Preprocessing

The acquired epifluorescence images were loaded into Python, then rolling ball subtraction was applied to the c-fos and DAPI channels using the scikit-image package with a rolling ball radius of approximately five times the size of a cell nucleus in the GCL or ML (to capture clusters of labeled nuclei in the granule cell layer). After background subtraction, a single threshold value for each channel was selected to capture the remaining signal while omitting background.

#### 4.6.2. Particle Analysis

To quantify fluorescently labeled particles, we used a custom Python-based pipeline to detect and count particles in the DAPI and c-Fos channels within manually defined ROIs surrounding the entire molecular layer. Following preprocessing, a Gaussian filter was applied to both channels with a sigma radius of 1.5. Next, the images were thresholded, and the corresponding ROI files were applied to create a mask excluding areas outside the region of interest. Distance transform-based watershed segmentation was then applied to delineate separations between individual particles, and particles were filtered based on size and circularity. The remaining detected particles in the DAPI channel were counted and used to create masks that were subsequently applied to the c-Fos channel. The number of c-Fos-positive particles contained within the DAPI masks was counted, and the final ratio of c-Fos-positive (active) to DAPI-positive (total) particles was computed. Output data included raw particle counts for each channel, computed ratios, and associated metadata, which were saved as .csv files for downstream statistical analysis. For quality control, visual outputs showing the raw image, ROI, threshold mask, and overlay were generated and saved.

#### 4.6.3. Fluorescent Area Analysis

Because the labeled nuclei in the granule cell layer of the cerebellar cortex were too dense to separate and count, a second analysis pipeline was applied to the c-Fos channel to quantify the relative area of fluorescent labeling within the entire granule cell layer of each lobule. Images were preprocessed and thresholded as described above, then the corresponding ROIs were applied to create masks excluding areas outside of the granule cell layer. The remaining thresholded area was computed and normalized to the total ROI area. Output data included raw area measures, the computed percent fluorescent area within the ROI (complete granule cell layer), and associated metadata, which were saved as .csv files for downstream statistical analysis. For quality control, visual outputs showing the raw image, ROI, threshold mask, and overlay were generated and saved.

### 4.7. Statistics

For each lobule, the cerebellar cortex was sampled across three adjacent 40 μm sections; fluorescence measurements from the same anatomical subregion across these sections were averaged to yield a single value per subregion per animal. Where the field of view at 20× magnification did not encompass a full lobule, spatially distinct subregions were imaged separately. A linear mixed model (LMM) with animal ID as a random intercept and Satterthwaite-approximated degrees of freedom was applied separately within each experimental group to test for statistically significant differences between subregions within a lobule. Pairwise comparisons were Bonferroni-corrected. Where no significant differences were detected, subregions were combined for subsequent group comparisons. Lobule 7 was the only region for which dorsal and ventral subregions were retained as distinct throughout. A second LMM was applied to test for laterality in cerebellar hemisphere data by comparing left and right hemisphere measurements within each lobule. No significant differences were detected in any lobule, and left and right hemisphere data were pooled for all subsequent analyses. Experimental group comparisons were subsequently performed for each lobule using LMMs with the same random effects structure and Bonferroni correction. All analyses were conducted in SPSS version 29.0.2.0.

## Author Contributions

Conceptualization, M.S., A.S.; methodology, M.S., V.Y; formal analysis, M.S., F.M.; data curation, M.S.; writing—original draft preparation, M.S.; writing—review and editing, A.S.; visualization, M.S.; supervision, A.S.; funding acquisition, A.S. All authors have read and agreed to the published version of the manuscript.

## Funding

A.S. received funding from the Canadian Institutes of Health Research (CIHR) Project Grant PJT-178281, Natural Sciences and Engineering Research Council of Canada (NSERC) Discovery Grant RGPIN-2020-07073, Canada Foundation for Innovation John R. Evans Leaders Fund (CFI-JELF) Equipment Grant 38053, Fonds de recherche du Québec – Santé (FRQS) Chercheurs Boursiers/Chercheuses Boursières, FRQS Établissement de jeunes chercheurs, a New Recruit Start-Up Supplement from Healthy Brains for Healthy Lives (HBHL) and the Canada First Research Excellence Fund (CFREF), and startup funding from the Research Institute of the McGill University Health Centre. M.S. received funding from FRQS, Faculty of Medicine Studentship and the McGill Integrated Program in Neuroscience, F.M received funding from HBHL.

## Conflicts of Interest

The authors declare no conflicts of interest.

## Notes

https://github.com/victor-yon/fluorescent-area-analysis/tree/main

### Competing Interest Statement

The authors have declared no competing interest.

### Summary of Updates

Title altered to match initial journal submission; scale bar descriptors added to figure captions.

